# Missing values are informative in label-free shotgun proteomics data: estimating the detection probability curve

**DOI:** 10.1101/2022.07.02.498573

**Authors:** Mengbo Li, Gordon K. Smyth

## Abstract

Mass spectrometry proteomics is a powerful tool in biomedical research but its usefulness is limited by the frequent occurrence of missing values in peptides that cannot be reliably quantified for particular samples. Many analysis strategies have been proposed for missing values where the discussion often focuses on distinguishing whether values are missing completely at random (MCAR), missing at random (MAR) or missing not at random (MNAR). We argue here that missing values should always be viewed as MNAR in label-free proteomics because physical missing value mechanisms cannot be identified for individual points and because the probability of detection is related to underlying intensity. We show that the probability of detection can be accurately modeled by a logit linear curve. The curve asymptotes close to 100%, limiting the potential role of missing values unrelated to intensity. The curve is also incompatible with simple censoring mechanisms. We propose a statistical method for estimating the detection probability curve as a function of the underlying intensity, whether observed or not. The model quantifies the bias of missing intensities as compared to those that are observed. The model demonstrates that missing values are informative and suggests possible approaches to imputation and differential expression.

## Introduction

Mass spectrometry (MS)-based shotgun proteomics is a powerful tool in biomedical research. Shotgun proteomics refers to the suite of bottom-up proteomics methods where proteins in complex biological samples are identified and quantified using liquid chromatography coupled with tandem mass spectrometry (LC-MS/MS) [1, 2]. Compared with label-based approaches, label-free quantification has gained its popularity as it involves less experimental preparation and has the capacity to process large numbers of samples [3, 4]. In a typical workflow, protein contents extracted from samples are first broken down into peptide mixtures through proteolytic digestion. Peptides are then separated by liquid chromatography and analysed by a mass spectrometer [5]. Peptide precursors are identified by correlating fragment ion spectra produced in the tandem mass spectrometry (MS/MS) step to the theoretical MS/MS spectra predicted in a protein sequence database or a spectral library [6]. Peptide abundances are measured by MS intensities derived from peak heights or areas. Protein quantification is then achieved by summarising intensities of constituent peptides. The past decade has seen advancements in many aspects of MS-based proteomics, both instrumental and methodological. Nevertheless, missing values are still common in label-free shotgun proteomics despite of the continuing technological upgrades. Consequently, care must be taken in data analysis where missing values often contain a substantial amount of information.

Rubin developed a framework for statistical analysis of datasets with missing values that has been highly influential [7, 8, 9, 10]. Rubin classified missing value processes into three broad categories that determine the type of statistical analysis that is possible and appropriate. The simplest case is when values are missing completely at random (MCAR) independently of the data values themselves. If MCAR applies, then the missing values can be ignored because the observed data are a simple random sample of the complete data. If the data are not MCAR but the missing value process depends only on observed covariates then statistical models can be fitted by conditioning on the observed data, for example using the widely popular EM algorithm [11]. This case is called missing at random (MAR). The remaining category is missing not at random (MNAR) when the missing value process depends on the underlying value itself. When values are MNAR, the missing values are non-ignorable and the data can only be analysed by estimating the relationship between the underlying data value and the probability that the value is missing [10, 12]. In this article, we say that a peptide or protein has been detected if the corresponding intensity value is observed and non-detected if it is not observed. We argue that only MNAR is relevant when designing analysis methods for MS proteomics data and we provide a method for estimating the probability of detection as a function of the underlying intensity value.

Rubin’s classification has often been misinterpreted [8]. The three MCAR, MAR and MNAR categories do not correspond to particular physical missing value mechanisms but rather they summarize the statistical information content of the missing values. Each dataset may be subject to multiple missing value mechanisms, some of which may be random. However, if the aggregate effect of the missing value mechanisms is such that the probability of detection is dependent on the underlying intensity, then the missing values are MNAR in terms of the statistical classification.

For proteomics data, the most likely missing value mechanisms are either (a) the expression level of the peptide precursor is below the detection limit of the instrument or (b) the elution profile or spectral signature of the precursor cannot be distinguished from the signatures of other precursors in the same sample [13]. The second mechanism is highly stochastic because it depends on the extent to which other precursors with interfering profiles are expressed in the same sample. Both mechanisms are intensity-dependent because the precursor’s signature is more likely to be identified if the peptide is more highly expressed relative to similar peptides in the same sample. The subtlety of proteomics data is that there is no clear-cut or fixed detection limit and even very highly expressed peptides can be subject to missing values, so it is difficult to assign specific causes to particular observations. The missing value process therefore needs to summarized probabilistically to be modeled accurately.

It is a common observation in proteomics publications that the frequency of missing values tends to decrease with peptide abundance [1, 14, 15, 16, 17]. In this paper we systematically explore the shape of this relationship. Luo et al [18] assumed the detection probability to be linearly dependent on its log-intensity on the logit scale. However, the proposed model has a complex hierarchical structure and is only suitable for iTRAQ data. O’Brien et al [19] assumed the detection probability to be probit-linear in the underlying log-intensity as part of a fully Bayesian differential expression approach. In this paper we propose a direct and generally applicable method for estimating the detection probability curve.

Our aim is to provide a formal statistical model for the detection probabilities in label-free shotgun proteomics data. Based on observed data, we empirically examine the relationship between the proportion of detected samples for each peptide precursor and the average log-intensity of that precursor. We show that the detection proportions increase with intensity. This increasing relationship is demonstrated on six public datasets of different numbers of detected peptides, different sample sizes, and different levels of missingness. We propose a statistical model that depicts the relationship between detection and intensity and which is visualized by a detection probability curve. The detection probability curve relates the probability of an observation being detected to its underlying intensity, whether or not that intensity was observed. The proposed model also provides an assessment of the biases caused by non-ignorable missing values in relative abundance estimation among samples, and hence has meaningful implications for the identification of differentially expressed proteins in the analysis of label-free shotgun proteomics data.

## Materials and methods

### Datasets

#### Dataset A: Hybrid proteome data

Hybrid proteome samples were generated by mixing human, yeast and E.coli lysates in different ratios and analysed by SWATH-MS. For each mixture, triplicates were measured [4]. The original data were published accompanying the LFQbench software [4] where details on sample preparation and data acquisition are available. We downloaded the HYE110 dataset (TripleTOF 6600; 64-variable-window acquisition) processed by DIA-NN under the library-based mode, with settings of DIA-NN detailed in Demichev et al [13]. Precursor-level intensities were extracted from the DIA-NN report. For our analysis, we only consider the triplicates of Sample A (*n* = 3). In this sub-sampled dataset, 34,689 precursor ions are detected in at least one sample, and the overall proportion of missing values is approximately 8.6%. We also set intensity values less than 1 to be missing; this affected only a small number of values and had little impact on the percentage of missing values or on the results presented here. The log-2 transformation was applied to the intensities.

#### Dataset B: Cell cycle proteomes

Single cell proteomes were profiled by the true single-cell-derived proteomics (T-SCP) pipeline as described in Brunner et al [20]. Four cell populations enriched in different cell cycle stages were produced from HeLa cells by drug treatment [20]. Precursor ions in prepared samples were fragmented in the parallel accumulation–serial fragmentation with data-independent acquisition (diaPASEF) mode [21]. The MS raw files were analysed by DIA-NN in library-based mode [13]. Details on sample preparation, LC-MS/MS analysis and data processing are provided in Brunner et al [20]. Processed data were downloaded from the ProteomeXchange Consortium via the PRIDE [22] partner repository with the dataset identifier PXD024043. Precursor-level output was obtained from the DIA-NN report and cells from the cell cycle experiment were extracted (*n* = 231). The number of detected precursor species is 10,754. About 60.4% of the data are missing values. We also set intensity values of zero to be missing; this affected only a small number of values and had little impact on the percentage of missing values. A log2-transformation was applied to the intensities.

#### Dataset C: HepG2 technical replicate data

Technical replicates of HepG2 cell lysates were analysed in Sinitcyn et al [23]. Briefly, MS data were collected by data-independent acquisition (DIA) and analysed by MaxDIA in discovery mode, a DIA data analysis software within MaxQuant [24]. Descriptions on sample preparation, LC-MS/MS procedures and the data processing workflow are described in Sinitcyn et al [23]. Processed data were downloaded from the ProteomeXchange Consortium via the PRIDE [22] partner repository with the dataset identifier PXD022589. Peptide-level data were obtained from the MaxQuant output, which reports missing values as zeros. Intensity values were log2-transformed. The number of detected precursors is 62,515 in 27 samples. The overall missing proportion is 32.8%.

#### Dataset D: Human blood plasma proteome

Human blood plasma samples from acute inflammation patients were collected and analysed by Prianichnikov et al [25]. In brief, MS data were acquired by data-dependent acquisition (DDA) using the timsTOF Pro mass spectrometer operated on the PASEF scan mode. Raw MS data were then analysed by MaxQuant. Details on sample preparation, LC-MS/MS procedures and the MaxQuant workflow are reported in Prianichnikov et al [25]. Processed data are available from the ProteomeXchange Consortium via the PRIDE [22] partner repository with the dataset identifier PXD014777. Peptide-level data were extracted from the MaxQuant output. Peptide species that had zero intensities in all samples were discarded. Intensity values were log2-transformed. The number of precursor species after filtering is 2,384 in 212 samples. The overall missingness proportion is 56.9%.

### Regression splines

Regression splines were fitted to the proportion of detected (non-missing) values for each precursor based on its average observed intensity. For peptide precursor *i*, write *p_i_* for the probability of detection. We modelled *p_i_* using logit regression splines of the form

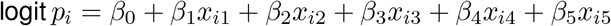

where the *x_ik_* are natural spline basis vectors computed by the ns() function in the splines package in R. The basis vectors were computed from the average observed log-intensities 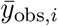 for each precursor. The number of basis vectors is called the degrees of freedom (df) for the spline. Splines were fitted with df= 1, 3 or 5. For df= 3 and df= 5 the spline knots were set according to the ns() function default, which separates the ordered 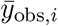 values into equal numbered groups. For df= 1, the logit-linear model:

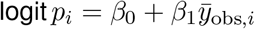

was used as the regression spline. When fitting regression splines at the protein-level data, basis vectors were generated on the average observed log-intensity for each protein (or protein group).

### Zero-truncated binomial distribution

The number of detected samples for each precursor was modeled by the zero-truncated binomial distribution, that is,

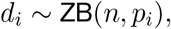

*n* being the sample size and *d_i_* being the number of detected intensity values in precursor *i*. The probability of detecting a sample is precursor-specific with *p_i_* ∈ [0,1]. The probability of detecting exactly *k* samples in precursor *i* is given by

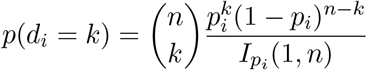

for *k* = 1,2,…,*n* and *I_pi_* (1,*n*) = 1 – (1 – *p_i_*)^*n*^. The spline regression parameters were estimated by maximum likelihood using the zero-truncated binomial distribution.

### Capped logit-linear model

The capped logit-linear model adds an asymptote parameter to the logit-linear model. The model assumes that the probability of detection for precursor *i* is

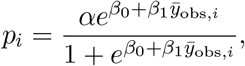

where 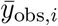 is the average observed log-intensity for that precursor. Here 0 < *α* ≤ 1 imposes a cap on the detection proportion for all precursors. Assuming that *β*_1_ > 0, *α* is the asymptotic probability of detection for precursors with high observed intensities. The capped logit-linear model was estimated by maximizing the zero-truncated binomial likelihood with respect to the parameters *β*_0_, *β*_1_ and *α*. The same model was applied also to protein-level intensities, in which case 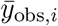 was the averaged observed log-intensity for protein *i*.

### Probability model for the missing value distribution

Let *y* be a log-intensity value and *d* be the indicator of detection with *d* = 1 if *y* is observed and *d* = 0 if *y* is missing. Write *f*_obs_(*y*) = *f*(*y*|*d* = 1) for the observed data distribution, i.e., the probability distribution for *y* conditional on *y* being observed. Similarly write *f*_mis_(*y*) = *f*(*y*|*d* = 0) for the missing data distribution, i.e., the probability distribution for y conditional on y being unobserved.

We assume that the detection probability is a logit-linear function of *y*,

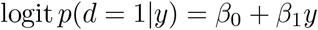

and that the observed data distribution is normal with mean *μ*_obs_ and variance 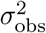. It can be proved that the missing data distribution is also normal with mean 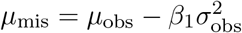 and variance 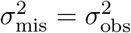. The marginal log-odds of detection can be shown to be

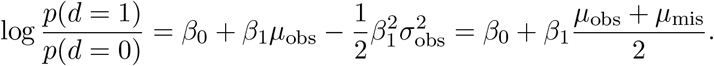

A complete mathematical derivation is given in Supplementary Methods Section 1.4.

## Results

### Detection increases with intensity in proteomics data

We first demonstrate the relationship between missingness and observed intensity on the precursor level using a variety of previously published proteomics datasets. We focus on datasets consisting of two or more replicate samples (biological or technical replicates) for two or more experimental conditions. Each dataset therefore consists of a matrix of log-intensities *y_ij_* for peptide precursors *i* = 1,…,*m* and samples *j* = 1,…,*n*. If a particular *y_ij_* is non-missing, then we say that precursor *i* is *detected* in sample *j*. We will write *d_ij_* = 1 if *y_ij_* is detected and *d_ij_* =0 if *y_ij_* is missing. We will further write 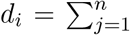 for the total number of detected (non-missing) values for precursor *i*.

Our aim is to explore the probability of detection as a function of the intensity value *y_ij_*, but we cannot do this directly because the intensities of non-detected precursors are unknown. Instead we take advantage of the fact that expression intensities typically vary by orders of magnitude across different peptide precursors but are relatively less variable across samples for the same precursor. For most datasets it is reasonable to assume that the majority of precursors are not differentially expressed between conditions. Even for differentially expressed precursors, the expression fold-changes are typically less than one order of magnitude with only a minority of larger fold-changes. We therefore examine the proportion of missing values for each precursor as a function of the mean of the observed log-intensities for that precursor, 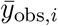, with the confidence that the observed log-intensities for each precursor are at least roughly representative of the likely magnitude of the unobserved intensities for the same precursor.

Figure 1 shows empirical logit spline curves fitted to the observed proportion of detected values for each peptide precursor for four public datasets. The proportion of detected samples increases as the average observed intensity increases on precursor-level data, regardless of the overall frequency of missing values in the dataset. The number of samples varies from *n* = 3 to *n* = 231 for the different datasets, but the increasing trend is discernible even for the smallest dataset. The same monotonic increasing relationship between average log-intensity and proportion detected was also observed when intensities were summarized at the protein level for the same datasets (Figure S1). Figures S5 and S6 show the same increasing relationship for two more datasets. This strong relationship between intensity and detection proportion shows that the missing values do not occur at random but are probabilistically related to peptide expression.

**Figure 1:**
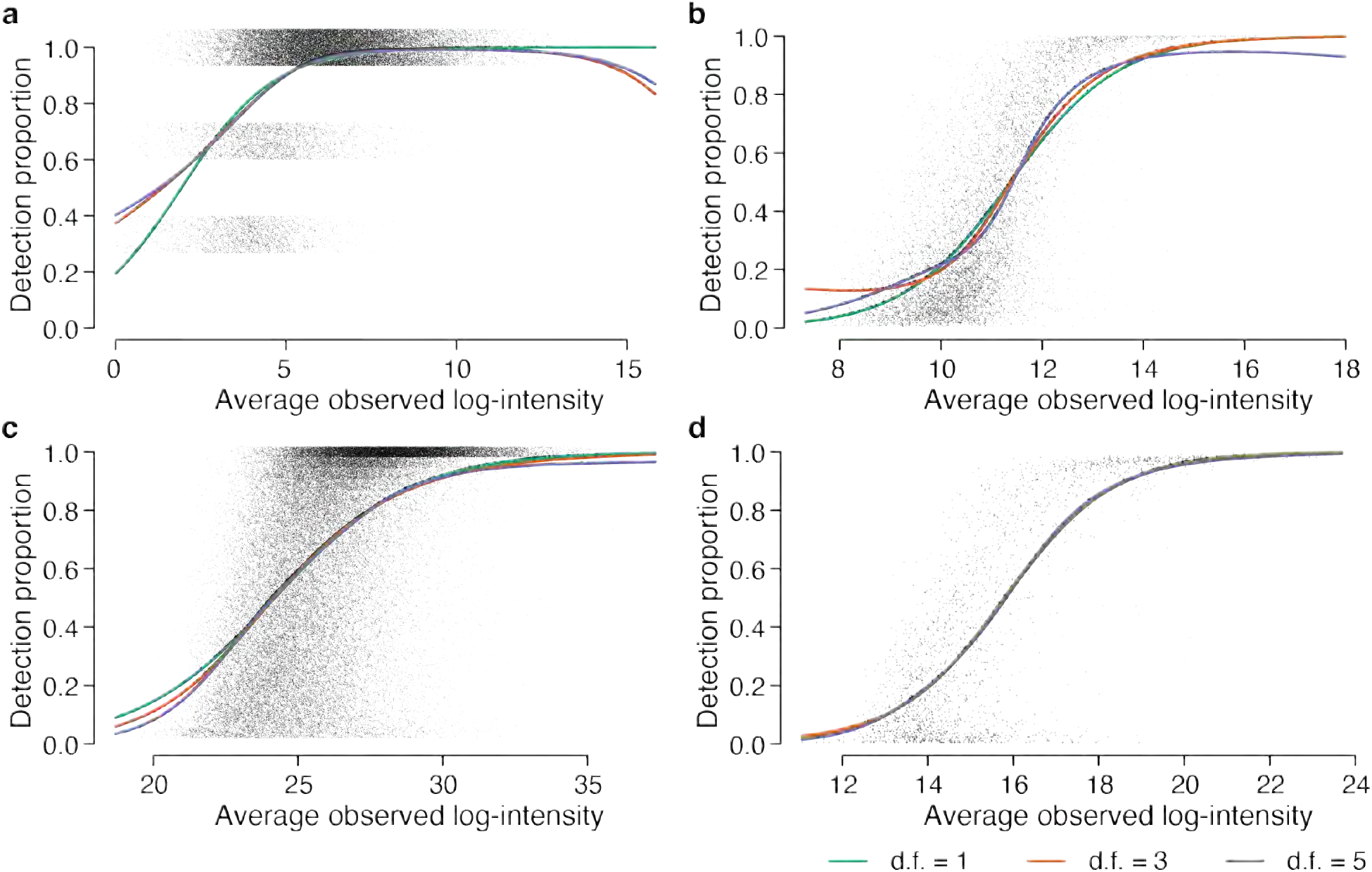
The proportion of detected values increases with the average intensity of each peptide precursor. Panels (a)–(d) show scatterplots for Datasets A–D. The x-axis shows average log2 observed intensity for each precursor. The y-axis shows the proportion of detected (non-missing) values for each precursor. The number of samples is (a) *n* = 3, (b) *n* = 231, (c) *n* = 27 and (d) *n* = 212. Jittering is added to the detection proportions in (a) and (c) to reduce over-plotting. Natural cubic splines were fitted to the logit proportions with 1, 3 or 5 degrees of freedom (df).

### The observed proportion of detected samples is zero-truncated

Precursors that are missing in all samples are omitted from Figure 1 because there are no observed intensities from which to compute 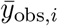. In Figure 1a, for example, the dataset contains *n* = 3 samples and only precursors with 1, 2 or 3 observed intensities can be shown on the plot. The absence of precursors with zero detected values needs to be taken into account when estimating the detection proportion curves. We model the number of detected samples for each precursor by a zero-truncated binomial distribution, i.e.,

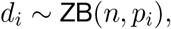

where *d_i_* is the number of detected samples for precursor *i, n* is the sample size, *p_i_* is the detection probability for that precursor and ZB denotes the zero-truncated binomial distribution. In Figure 1, we model logit(*p_i_*) as a spline function of the average log-intensity *y*_obs,*i*_ and the parameters of the spline are estimated by maximum likelihood using the truncated binomial likelihood.

To demonstrate the effect of using the zero-truncated binomial distribution, we re-fitted the spline curves to Dataset A using both zero-truncated and standard binomial distributions. The detection probability is consistently overestimated at lower intensities if the ordinary binomial distribution is used (Figure 2). The overestimation remains regardless of the number of parameters used for the regression spline. The zero-truncation adjustment provides accurate estimation of detection probabilities for lower intensity precursors especially when the sample size is limited.

**Figure 2:**
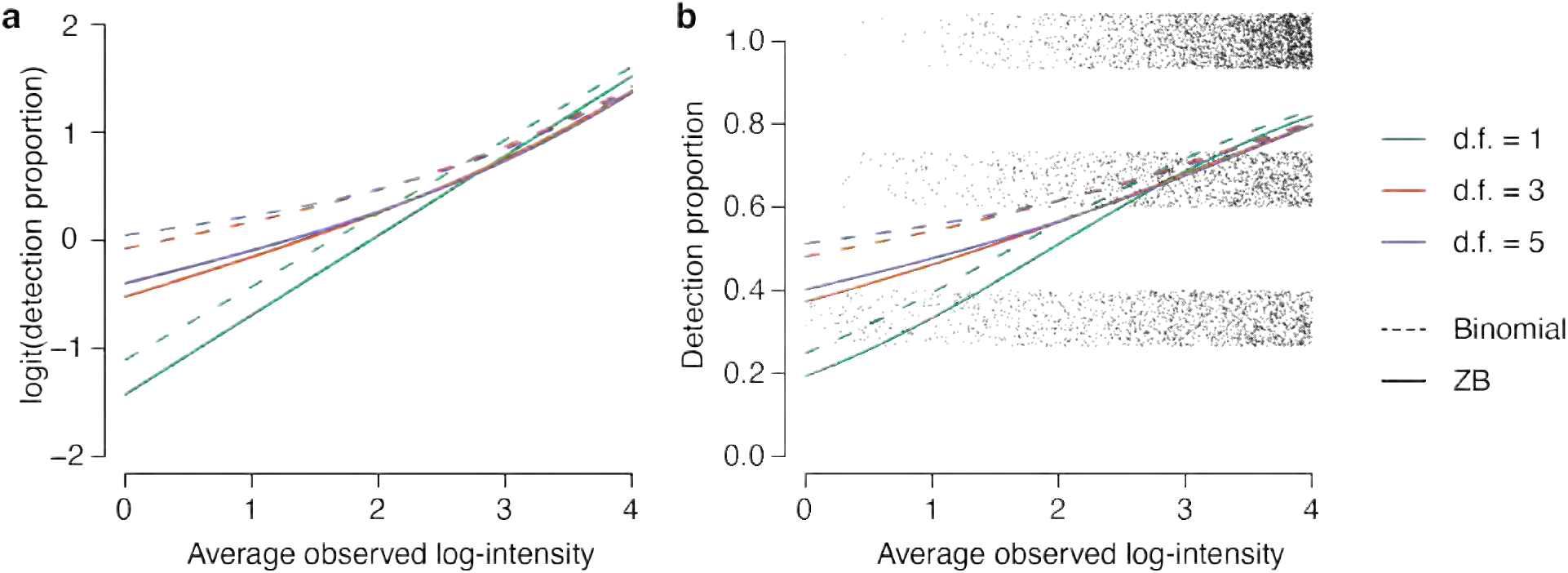
Detection probabilities are biased if zero-truncation is ignored. Nature regression splines are fitted to the Dataset A logit detection proportions by maximizing either the ordinary binomial likelihood (dashes lines) or the zero-truncated binomial likelihood (solid lines). The splines use either 1, 3 or 5 df (green, red and black respectively). Panel (a) plots detection proportions on the logit scale and panel (b) shows untransformed proportions. Jittering is added to the detection proportions in (b) to reduce over-plotting. The detection probabilities are consistently over-estimated by the ordinary binomial likelihood.

### The detection proportion is approximately logit-linear in log-intensity

The spline curves in Figure 1 were estimated with either 1, 3 or 5 degrees of freedom (df), with the df being equal to the number of regression coefficients estimated in the fit. For each dataset, the three fitted curves are not materially different, suggesting that the logit-linear curve, with 1 degree of freedom, is sufficient to summarize the intensity-dependent trend. To explore this more rigorously, we computed the percentage of deviance explained by each regression spline coefficient. We defined the total deviance to be twice the log-likelihood difference between most complex spline regression model with 5 df and the null model with only an intercept term. The deviance explained by each spline degree of freedom can then be evaluated by increases in the log-likelihood as parameters are added to the spline curve. We found that almost all of the log-likelihood difference was explained by the linear coefficient (Table 1). The logit-linear coefficient explains over 96% of the total deviance for all datasets. The four non-linear parameters together explain less than 4% of the deviance. At the protein-level, the logit-linear trend explains over 97% of the deviance (Table S2). Similar results were observed for datasets E and F (Figure S4c, S5c). These results suggest that the probability of detection for each precursor (or protein) is approximately linear in log-intensity on the logit scale. We will therefore assume that the detection probability can be represented as a logit-linear function of log-intensity for the remainder of this article.

**Table 1:** Percentage of total deviance explained by the logit-linear and non-linear regression splines in Figure 1. The rows of the table show the percentage of the total deviance explained by the the logit-linear curve and the additional deviance explained by logit splines with 3 or 5 df. The second row gives the additional deviance explained by df= 3 over df= 1 and the third row the additional deviance explained by df= 5 over df= 3. The logit-linear curve explains > 96% of the deviance for all datasets.

| Source | Deviance Explained (%) |  |  |  |
| --- | --- | --- | --- | --- |
|  | A | B | C | D |
| Linear | 97.242 | 96.40 | 99.438 | 99.896 |
| Non-linear 2–3 | 2.732 | 2.32 | 0.328 | 0.029 |
| Non-linear 4–5 | 0.026 | 1.28 | 0.234 | 0.075 |

### 0.1 Missingness is negligible for high intensity precursors

The detection proportion curves shown in Figure 1 approach 100% for high observed intensities. If some of the missing values occur for reasons that are unrelated to the intensity level, then a proportion of missing values should persist even for very high intensity precursors. To explore whether this is true, we extended the logit-linear model to allow the curve to asymptote at a value less than 100%. The detection probability curves were found to asymptote at values very close to 1 for all datasets (Figure 3). The residual proportion of missing values for large intensities is always less than 3% and usually less than 1%. This shows that missing value mechanisms that are unrelated to intensity must be limited to a very small proportion of precursors. The same phenomenon is observed for intensities summarized at the protein-level (Figure S2) and for Supplementary Datasets E and F (Figures S4 and S5).

**Figure 3:**
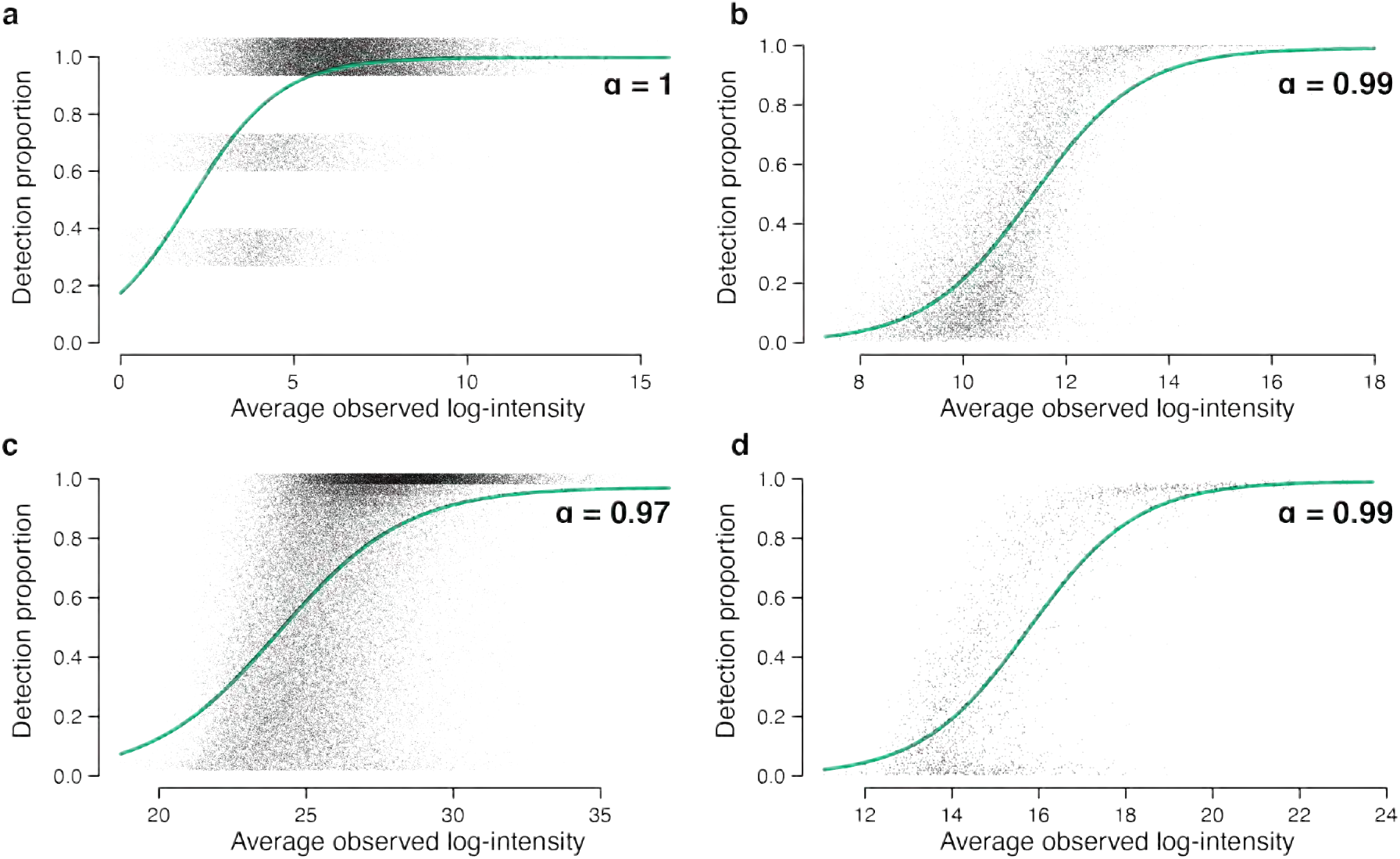
Capped logistic-linear curves for detection proportion. A logistic linear model is fitted to detection proportion on average observed intensity for Datasets A-D in (a)-(d) on the precursor level. Detection proportions for all observations are capped by *α*, where *α* ∈ (0,1]. The maximum likelihood estimates for the asymptotic probability (*α*) are (a) 0.9974, (b) 0.9915, (c) 0.9713 and (d) 0.9912. Jittering is added to vertical axes of (a) and (c) to reduce over-plotting in precursors.

### The detection probability curve models non-ignorable missingness in proteomics data

So far we have examined empirically the relationship between missingness and intensity, leveraging the observed data. We now propose a formal model for the detection probabilities that allows us to estimate the probability that an observation is detected (non-missing) given its own underlying intensity, even when that underlying intensity is not observed.

As before, let *y_ij_* be a log-intensity value and *d_ij_* indicate detection with *d_ij_* = 1 if *y_ij_* is observed and *d_ij_* =0 if *y_ij_* is missing. If *d_ij_* = 0, then *y_ij_* represents the intensity that would have been returned by peptide quantification if the missingness mechanism had been absent or had not operated. In particular, each missing *y_ij_* reflects the true expression level of that precursor in that sample in the same way that observed intensities do.

We assume a logit-linear relationship between the detection probability and the underlying intensity,

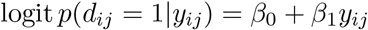

where *p*(*d_ij_* = 1|*y_ij_*) is the probability of detection conditional on *y_ij_*. We expect *β*_1_ > 0 as higher intensity values are more likely to be detected. The detection probabilities are precursor- and sample-specific, but the coefficients *β*_0_ and *β*_1_ are assumed to be appropriate across all precursors and samples. For simplicity of notation, we will drop the subscripts *ij* for the rest of this section, although our discussion is still for a specific precursor and sample.

In practice we only observe *y* when *d* = 1 and it is common to treat the observed values as normally distributed. We therefore assume that, conditional on being observed, *y*|*d* = 1 follows a normal distribution with mean *μ*_obs_ and variance 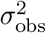. The key question is, what can be said about the unobserved value when *d* = 0? Under the above assumptions, it can be shown (see Supplementary Methods) that, when unobserved, the intensity *y*|*d* = 0 also follows a normal distribution with the same variance but a decreased mean. Specially, *y*|*d* = 0 follows a normal distribution with variance 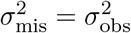 and mean

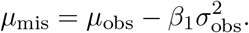

This reveals that *β*_1_*σ*_obs_ represents the number of standard deviations by which the missing values are biased relative to the observed values.

For estimation purposes, we need the marginal detection probability, not conditional on a possibly unobserved intensity value. It can be shown (Supplementary Methods), that the marginal log-odds of detection can be written as

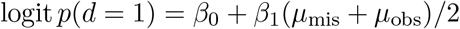

or equivalently

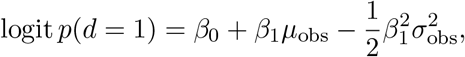

and this defines the detection probability curve. The probability of detection depends only on the global co-efficients *β*_0_ and *β*_1_ and on precursor-wise quantities *μ*_obs_ and 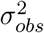 that can be estimated from the observed intensities.

To estimate *β*_0_ and *β*_1_, we take advantage of the fact that, for a well designed experiment, most precursors will not be differentially expressed between experimental conditions. It is therefore reasonable to assume that *μ*_obs_, 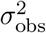 and *p*(*d* = 1) are precursor-specific but not sample-specific. We can therefore substitute in the observed precursor means and variances for *μ*_obs_ and 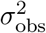 and then estimate *β*_0_ and *β*_1_ from the precursor-level detection proportions.

Figure 4 shows the detection probability curves estimated for Datasets A–D. For each precursor, *μ*_obs_ is estimated by the average observed log-intensity and the variance parameter 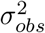 is estimated as the empirical Bayes moderated sample variance. Logistic regression coefficients *β*_0_ and *β*_1_ are estimated by maximum likelihood estimation assuming the zero-truncated binomial distribution for counts of observed samples in each precursor. The x-axis is (*μ*_mis_ + *μ*_obs_)/2, which represents the precursor mean log-intensity with observed and missing intensities equally weighted. The estimated *β*_0_ and *β*_1_ parameter values are given in Table 2. The detection probability curve generally reaches each detection probability at a slightly lower log-intensity once underlying intensities are taken into account, because the observed intensities are biased slightly towards higher values for each peptide precursor.

**Figure 4:**
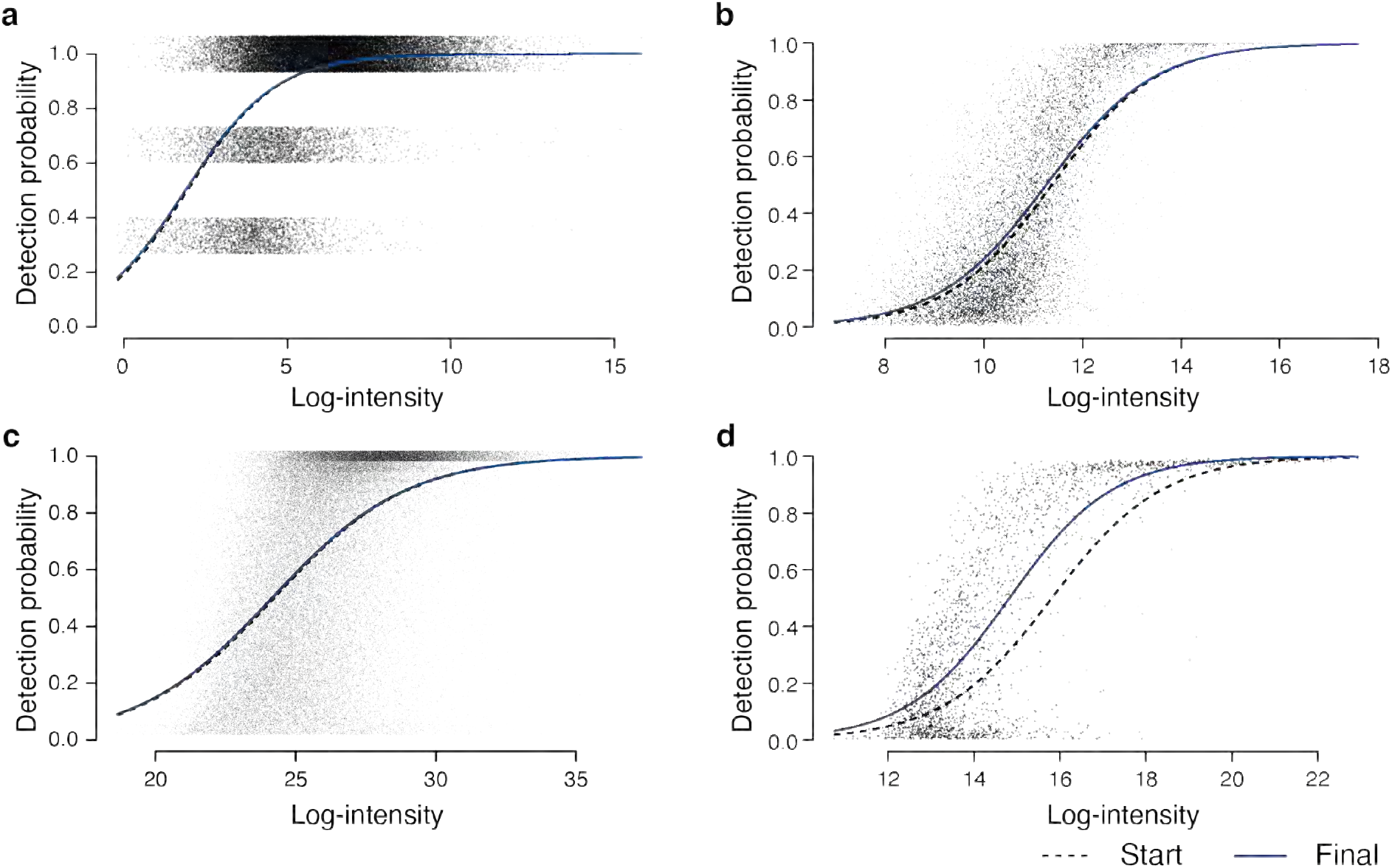
Detection probability curves. Panels (a)–(d) show estimated detection probability curves for Datasets A-D at the precursor level. In each dataset, the starting curve is obtained by fitting a logistic linear curve for detection proportions to average observed intensities whereas the final curve relates detection probabilities to underlying log-intensities. Jittering is added to vertical axes in (a) and (c) to reduce overplotting in precursors. The x-axis here is (*μ*_mis_ + *μ*_obs_)/2, which is the precursor mean log-intensity with observed and missing intensities equally weighted. The estimated curve parameters are given in Table 2.

**Table 2:** Parameter estimates for detection probability curves fitted on Datasets A–D that are visualized in Figure 4. The lower (Underlying) values are the final estimates that related probability of detection to underlying log-intensity. The upper (Observed) values are the more naive estimates that are obtained by fitting a logit-linear trend to the observed log-intensities for each precursor.

| Detection probability curve |  |  |  |
| --- | --- | --- | --- |
| Dataset | Fitted on | $\beta_0$ | $\beta_1$ |
| A | Observed | -1.4264 | 0.7363 |
|  | Underlying | -1.3612 | 0.7258 |
| B | Observed | -10.7271 | 0.9430 |
|  | Underlying | -10.3113 | 0.9154 |
| C | Observed | -10.1931 | 0.4212 |
|  | Underlying | -10.0860 | 0.4176 |
| D | Observed | -12.4087 | 0.7847 |
|  | Underlying | -12.4435 | 0.8393 |

## 1 DISCUSSION

Many previous articles have observed that lower intensity peptides tend to yield more missing values [1, 14, 16, 15]. Here we demonstrate that this is a universal phenomenon for label-free shotgun proteomics. The trend is seen for both protein- and precursor-level data and for datasets with high or low overall rates of missingness. We show that the detection probability can be accurately modeled as a logit-linear function of the underlying log-intensity. We point out the importance of the zero-truncated binomial distribution to avoid underestimating the rate of missing values. We show that the proportion of missing values becomes negligible for peptide precursors or proteins with sufficiently high intensities. We introduce the concept of a detection probability curve that relates the probability of detection to the, possibly unobserved, underling intensity. We propose a mathematical model that allows the detection probability curve to be estimated and which quantifies the relative bias between observed and unobserved intensities. The detection probability curve elicits the underlying relationship between missingness and intensity in a proteomics experiment and provides access to the hidden information contained in missing values. Our work is related to the Bayesian model proposed by O’Brien et al [19], but our approach here allows the detection probability curve to be explicitly estimated and explored.

There has been much discussion in the literature about the relevance of Donald Rubin’s MAR, MCAR and MNAR classification for proteomics data [26, 15, 16, 27, 28, 17, 29, 30]. Our analysis shows unambiguously that intensity is a strong predictor of the rate of missingness. This relationship is the key characteristic of MNAR and makes MAR and MCAR analyses generally unrealistic. It is important to understand that Rubin’s classification is statistical in nature in that the classification relates to the information available to the analyst rather than to physical mechanisms. We cannot rule out special missing values mechanisms that occasionally operate at random or which depend on factors other than the underlying intensity. However, if we do not know which missing values correspond to which mechanisms, then only the overall relationship between detection and intensity is relevant to the analyst. Since such detailed observation-specific mechanism information is never available to the analyst, it follows that all missing values are MNAR from a statistical point of view. Our results show that missing value mechanisms unrelated to intensity must be very rare if they do operate at all but, even if they were more common, a statistical analysis based on MNAR assumptions would still be appropriate.

The most common MNAR model is to assume that missing values are left-censored [14, 31, 26, 32], which is equivalent to a detection probability curve that is a step function. Our work shows that a more gradual detection probability curve is a better representation of missing value patterns.

The approach outlined here has implications for missing value imputation and for differential expression analyses of proteomics data. Our model allows the distribution of the unobserved intensities to be estimated from the observed values and a possible future approach would be to impute the missing values from this estimated distribution. The probabilistic approach allows the possibility of incorporating peptide-speciffc information into the missing value distribution so that imputation may be peptide-speciffc. The challenge in these or similar approaches will be to correctly propagate the uncertainty of estimation and uncertainty of imputation into the downstream differential expression analysis.

## Conclusion

Missing values in MS-based proteomics data are informative regarding the expression level of the peptide or protein and should be treated as occuring at random. The detection probability curve can be estimated as a function of the underlying intensity, whether observed or not. The detection probability curve provides a summary of the relationship between expression and missingness that clarifies the bias of missing intensities as compared to those that are observed.

## Data availability

The methods presented in this article have been implemented in the protDP package for R. Data and code to reproduce the results shown in this article are available from https://mengbo-li.github.io/protDP/. All datasets analysed in this article are publicly available as described in Methods.

## Supplementary data

Supplementary Data are available in the file supp.pdf.

## Acknowledgement

Thanks for Pedro Baldoni for helpful discussion and comments on an earlier version of the manuscript.

