## Supplementary Methods, Figures and Methods for "Missing values are informative in label-free shotgun proteomics data: estimating the detection probability curve"

### 1 Supplementary Methods

#### 1.1 Protein-level intensities for Datasets A-D

The missingness-intensity relationship was also examined for Datasets A-D on the protein group level. For the two DIA-NN [1] processed datasets, that is, Datasets A and B, MaxLFQ [2] intensities were obtained from their respective DIA-NN reports. For the two MaxQuant processed datasets [3], i.e., Datasets C and D, the `proteinGroups.txt` tables output by the MaxQuant software were used as the protein group level summary. The overall proportions of missing data are respectively $5.0\%$, $47.4\%$, $5.5\%$ and $41.2\%$ in Datasets A, B, C and D. The numbers of protein groups detected in at least one sample from the corresponding dataset are respectively $5,974$; $2,661$; $6,282$ and $332$ for Datasets A-D.

#### 1.2 Dataset E: Sydney Heart Bank

Cryopreserved left ventricular myocardium samples were analysed from donor hearts procured but not used for heart transplantation [4]. Mass spectrometry (MS) data were acquired in data-independent acquisition (DIA) mode and analysed by Spectronaunt v12, using a spectral library generated from fractions of the pooled mixture of all samples and analysed by LC-MS/MS in data-dependent acquisition (DDA) mode. Details on sample preparation, LC-MS/MS experiment workflow and data processing are available in [4]. Here we consider the healthy donor heart samples. Both precursor- and protein group-level data are log2-transformed before analysis. For the precursor-level analysis, there are $42,742$ precursors detected in at least one of the $24$ samples with an overall missingness proportion of approximately $33.4\%$. To obtain protein-level intensities, MaxLFQ [2] was applied. In protein-level data, the proportion of missing data is about $12.0\%$ in $3,208$ proteins. The dataset is publicly available from the ProteomeXchange Consortium via the PRIDE [5] partner repository with the dataset identifier PXD018678.

#### 1.3 Dataset F: UPS1 spiked-in yeast extract

Three concentrations of UPS1 (25 fmol, 10 fmol and 5 fmol) were spiked in yeast extract in triplicates [6]. MS data were obtained in DDA mode and processed by MaxQuant [3]. Here we look at the dataset that compares 25 fmol to 10 fmol spiked-ins. Processed data were downloaded from ProteomeXchange Consortium via the PRIDE [5] partner repository with the dataset identifier PXD002370. For peptide-level data, we used the `peptides.txt` file and for the protein-level analysis, we used the `proteinGroups.txt` file from the MaxQuant output. Log2-transformation was applied to precursor- and protein group-level intensities. In the peptide-level data, we observe $13,186$ peptide species detected in at least one of the $6$ samples. The missingness proportion is $14.7\%$. In the protein group-level data, we observe an overall proportion of $6.8\%$ missing values and $2,342$ protein groups.

#### 1.4 Mathematical derivation of detection probability curve

##### 1.4.1 Observed and missing distributions

Let $y$ be a log-intensity value and $d$ be the indicator of detection with $d = 1$ if $y$ is observed and $d = 0$ if $y$ is missing. Write $f_{\text{obs}}(y) = f(y|d = 1)$ for the observed data distribution, i.e., the probability distribution for $y$ conditional on $y$ being observed. Similarly write $f_{\text{mis}}(y) = f(y|d = 0)$ for the missing data distribution, i.e., the probability distribution for $y$ conditional on $y$ being unobserved.

It follows from Bayes theorem that

$$f_{\mathrm{obs}}(y) = f(y|d=1) = \frac{p(d=1|y)f(y)}{p(d=1)}$$

and

$$f_{\mathrm{mis}}(y) = f(y|d=0) = \frac{p(d=0|y)f(y)}{p(d=0)}.$$

where $f(y)$ is the marginal density distribution of $y$, $p(d=1|y)$ is conditional detection probability, $p(d=1)$ is the marginal detection probability and $p(d=0) = 1 - p(d=1)$. The ratio of the missing to observed density functions is therefore related to the detection probabilities by

$$\frac{f_{\mathrm{mis}}(y)}{f_{\mathrm{obs}}(y)} = \frac{p(d=0|y)}{p(d=1|y)}\frac{p(d=1)}{f(d=0)}.$$

##### 1.4.2 Assume logit-linear detection probabilities

We assume that the detection probability is a logit-linear function of $y$,

$$\operatorname{logit} p(d=1|y) = \beta_0 + \beta_1 y.$$

It follows that

$$\frac{f_{\mathrm{mis}}(y)}{f_{\mathrm{obs}}(y)} = \exp(-\beta_0 - \beta_1 y)\frac{p(d=1)}{p(d=0)}$$

which implies the missing value density is related to the observed value density by

$$f_{\mathrm{mis}}(y) = \frac{p(d=1)}{p(d=0)}\exp(-\beta_0 - \beta_1 y)f_{\mathrm{obs}}(y)$$

We know that the right-hand-side must integrate to 1. Also by definition

$$\int \exp(-\beta_1 y)f_{\mathrm{obs}}(y)dy = M_{\mathrm{obs}}(-\beta_1)$$

where $M_{\mathrm{obs}}()$ is the moment-generating function of the observed distribution. Hence we can conclude that

$$M_{\mathrm{obs}}(-\beta_1) = \exp(\beta_0)\frac{p(d=0)}{p(d=1)}$$

and therefore

$$f_{\mathrm{mis}}(y) = \frac{e^{-\beta_1 y}}{M_{\mathrm{obs}}(-\beta_1)}f_{\mathrm{obs}}(y).$$

##### 1.4.3 Assume observed values are normal

Let us assume now that the observed values follow a normal distribution, i.e., $f_{\mathrm{obs}}(y)$ is a normal density with mean $\mu_{\mathrm{obs}}$ and variance $\sigma^2_{\mathrm{obs}}$. The normal moment generating function is

$$M_{\mathrm{obs}}(-\beta_1) = \exp(-\beta_1\mu_{\mathrm{obs}} + \frac{1}{2}\beta_1^2\sigma^2_{\mathrm{obs}})$$

so

$$f_{\mathrm{mis}}(y) = \exp\left(\beta_1\mu_{\mathrm{obs}} - \frac{1}{2}\beta_1^2\sigma^2_{\mathrm{obs}} - \beta_1 y\right)\left(2\pi\sigma^2_{\mathrm{obs}}\right)^{-1/2}\exp\left\{-\frac{(y-\mu_{\mathrm{obs}})^2}{2\sigma^2_{\mathrm{obs}}}\right\}$$

which is the density of a normal distribution with mean

$$\mu_{\mathrm{mis}} = \mu_{\mathrm{obs}} - \beta_1\sigma^2_{\mathrm{obs}}$$

and variance

$$\sigma^2_{\mathrm{mis}} = \sigma^2_{\mathrm{obs}}.$$

##### 1.4.4 Marginal log-odds of detection

The marginal log-odds of detection can be written as

$$\begin{aligned}\log \frac{p(d=1)}{p(d=0)} &= \log \frac{\exp(\beta_0)}{M_{\text{obs}}(-\beta_1)} \\ &= \beta_0 + \beta_1 \mu_{\text{obs}} - \frac{1}{2}\beta_1^2 \sigma_{\text{obs}}^2 \\ &= \beta_0 + \beta_1 \frac{\mu_{\text{obs}} + \mu_{\text{mis}}}{2}.\end{aligned}$$

#### 2 Supplementary Figures

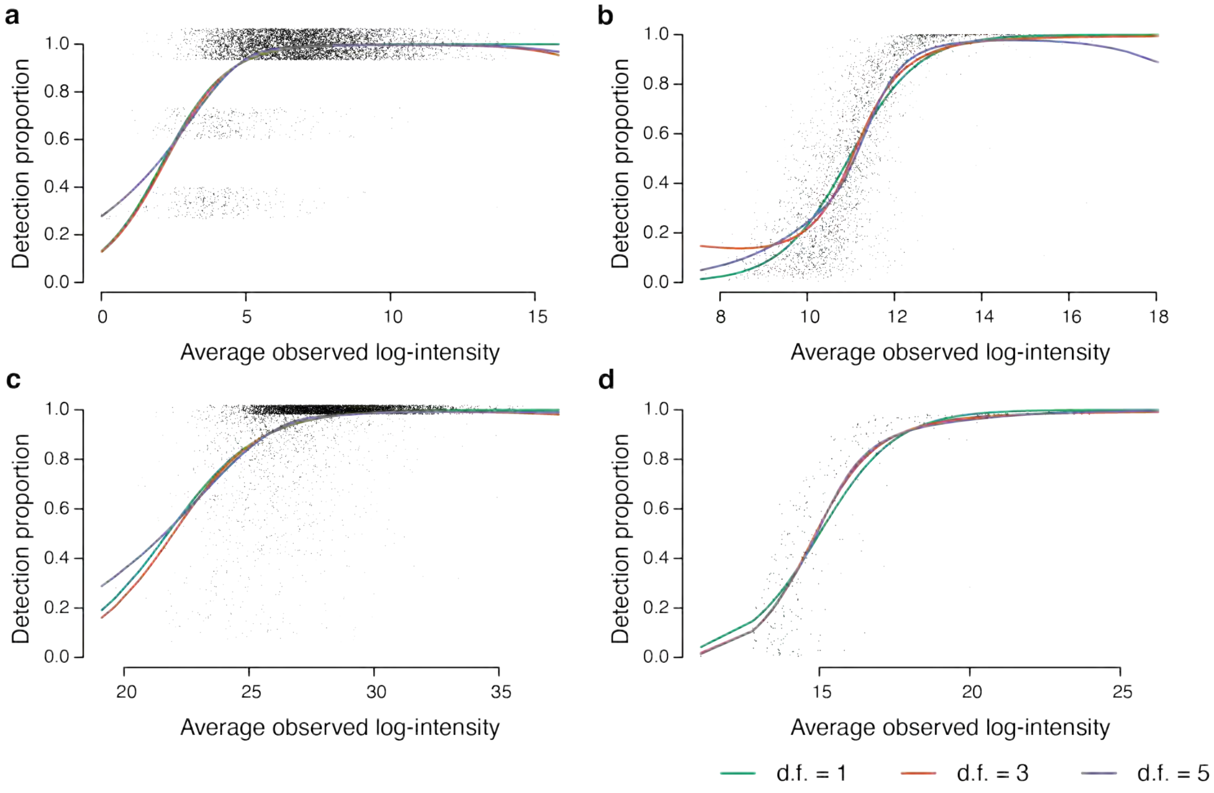


**Figure S1:** The proportion of detected samples increases in average intensity in each protein group. The x-axis shows the average observed intensity and the y-axis shows the proportion of detected samples in each protein group. Regression splines are fitted on the protein group-level for Datasets A–D in panels (a)–(d). Jittering is added to detection proportions in (a) and (c) to reduce over-plotting.

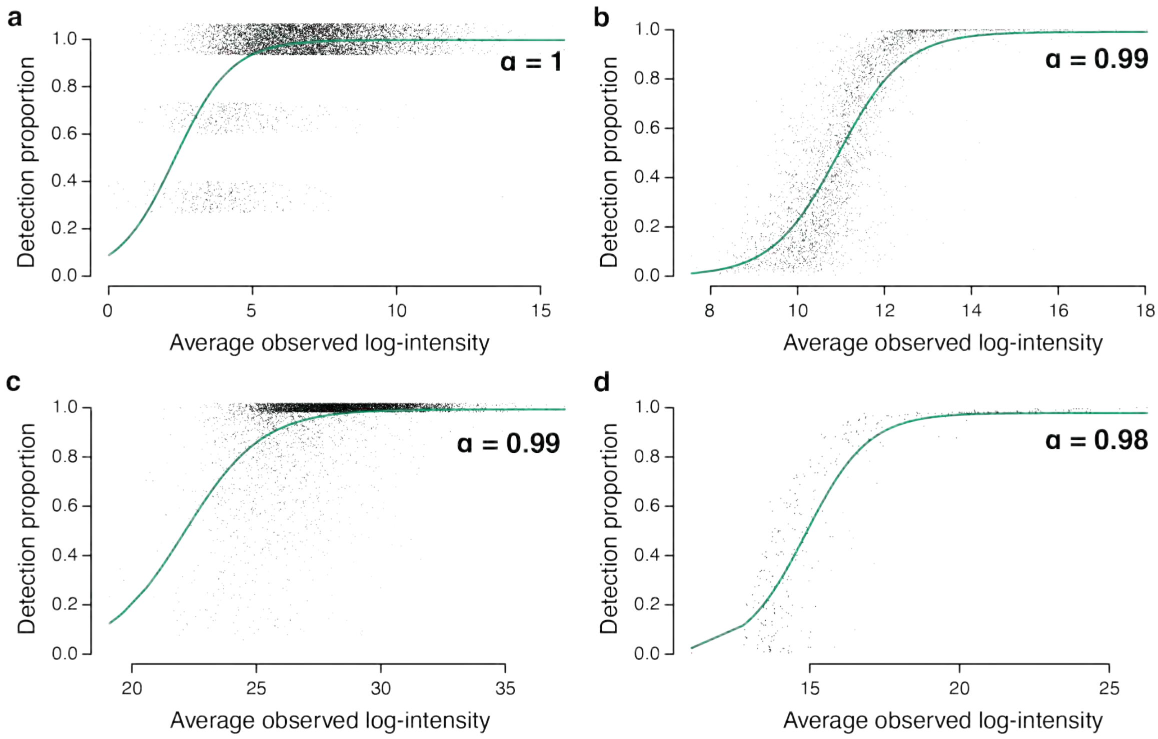


**Figure S2:** Capped logistic-linear curves for detection proportion on the protein group level. A logistic linear model is fitted to detection proportion on average observed intensity in each protein group for the Datasets A–D in (a)–(d). Detection proportions for all observations in the dataset are capped at $\alpha \in (0, 1]$. The maximum likelihood estimate of $\alpha$ for each dataset is indicated in (a)–(d). Jittering is added to vertical axes of (a) and (c) to reduce over-plotting in protein groups.

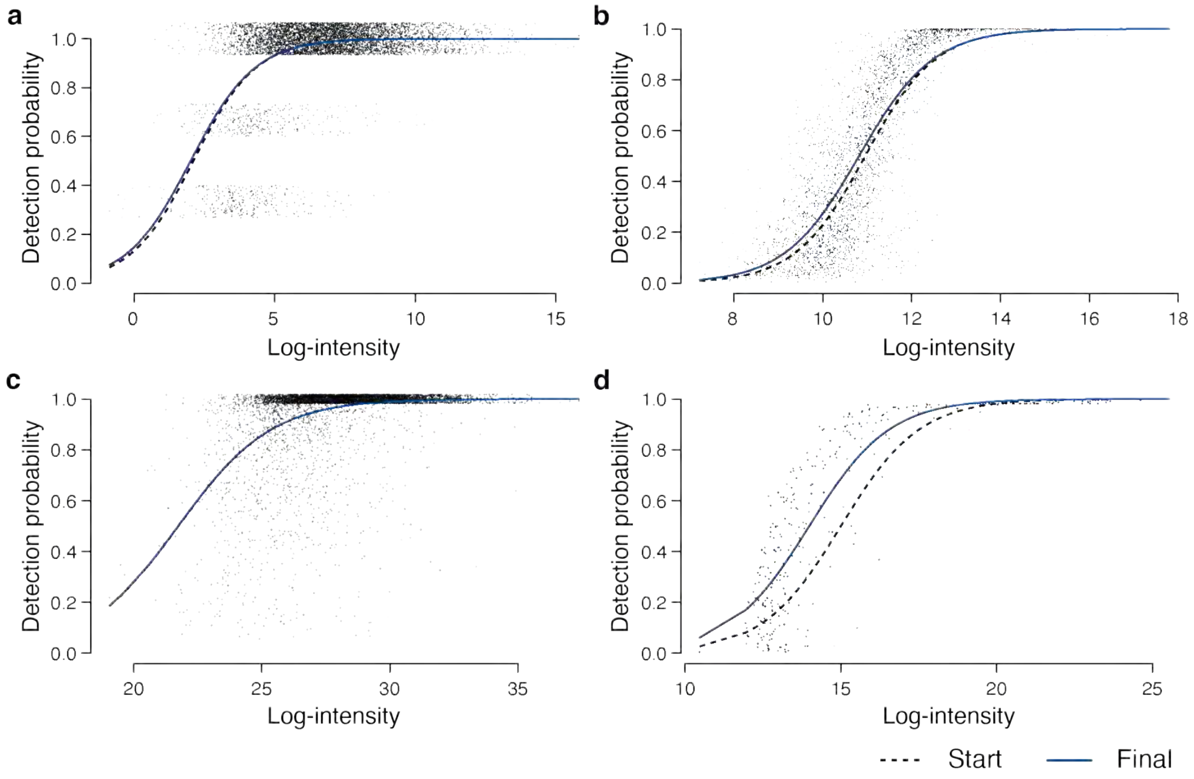


**Figure S3:** Detection probability curves fitted on the protein group level for Datasets A–D in (a)–(d). The starting curve is obtained by fitting a logistic linear curve for detection proportions to the average observed intensities on the protein group level. Jittering is added to vertical axes in (a) and (c) to reduce over-plotting. The estimated parameters for each curve are displayed in Table S2.

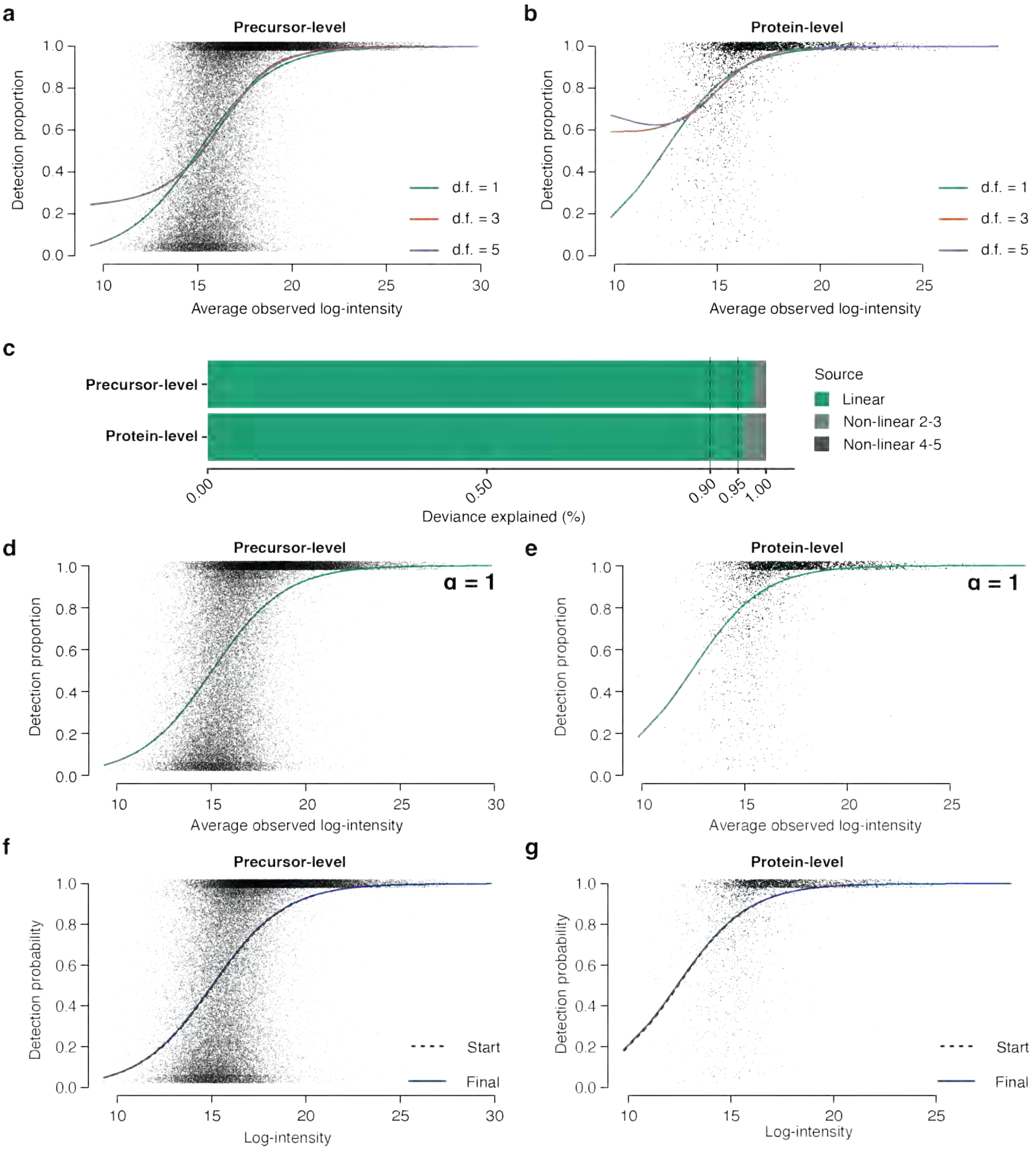


**Figure S4:** Relationship between missingness and intensity in Sydney Heart Bank data (Dataset E) on both precursor and protein group levels. (a) Regression splines fitted for detection proportions to average observed intensities in precursors. (b) Regression splines fitted for detection proportions to average observed intensities in protein groups. (c) Percentage of total reduced deviance explained by the logit-linear and non-linear regression splines from panels a and b. (d) Capped logistic-linear curve fitted for detection proportions to average observed intensities in precursors. (e) Capped logistic-linear curve fitted for detection proportions to average observed intensities in protein groups. (f) Detection probability curve fitted on the precursor level. (g) Detection probability curve fitted on the protein group level.

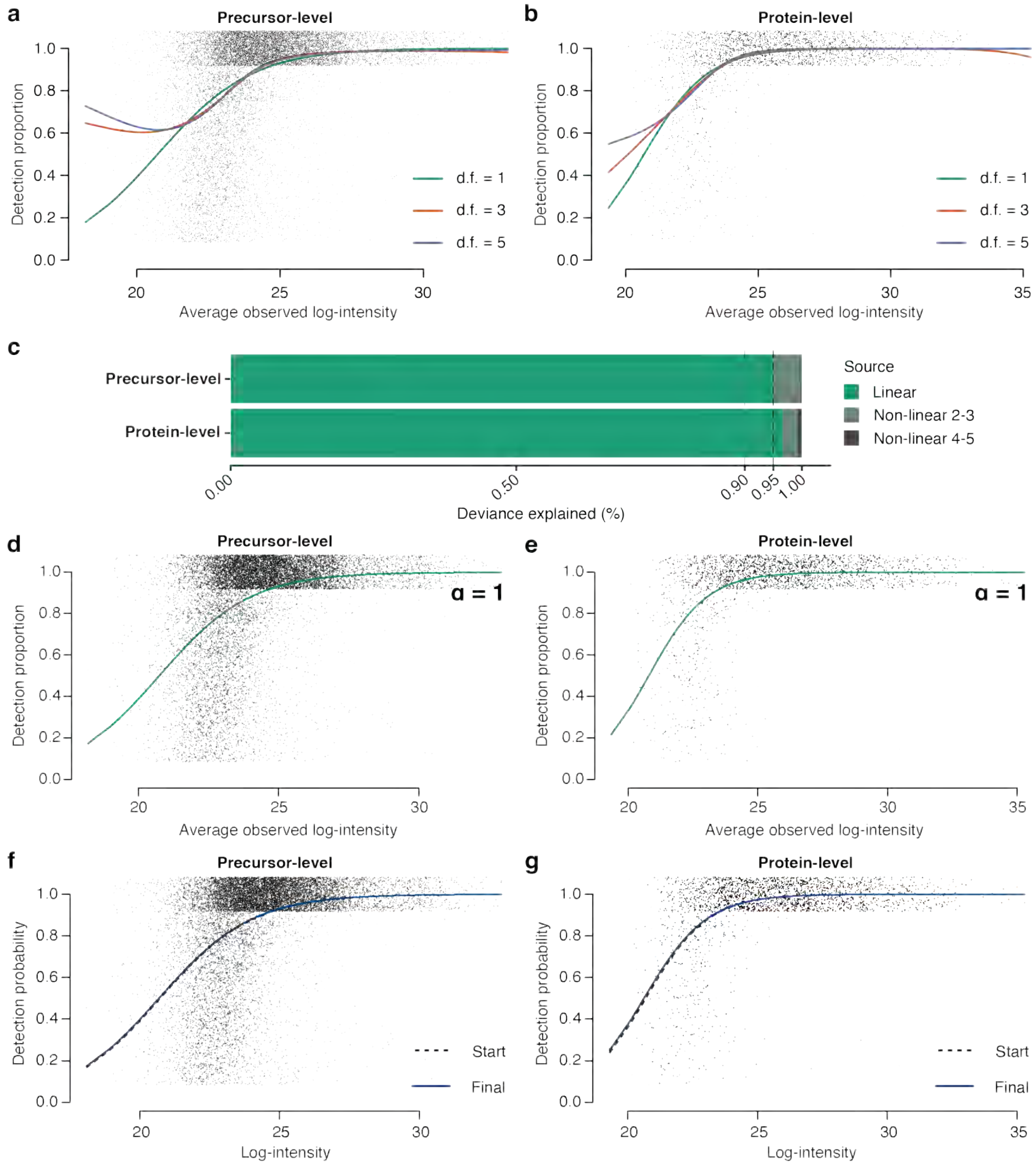


**Figure S5:** Relationship between missingness and intensity in UPS1 spiked-in yeast extract data (Dataset F) on both precursor and protein group levels. (a) Regression splines fitted for detection proportions to average observed intensities in precursors. (b) Regression splines fitted for detection proportions to average observed intensities in protein groups. (c) Percentage of total reduced deviance explained by the logit-linear and non-linear regression splines from panels a and b. (d) Capped logistic-linear curve fitted for detection proportions to average observed intensities in precursors. (e) Capped logistic-linear curve fitted for detection proportions to average observed intensities in protein groups. (f) Detection probability curve fitted on the precursor level. (g) Detection probability curve fitted on the protein group level.

#### 3 Supplementary Tables

**Table S1:** Percentage of total reduced deviance explained by the logit-linear and non-linear regression splines fitted on the protein group-level data shown in Figure S1. The rows of the table show the percentage of the total deviance explained by the the logit-linear curve and the additional deviance explained by logit splines with 3 or 5 degrees of freedom.

| | Deviance Explained (%) | | | |
|---|---|---|---|---|
| **Source** | **A** | **B** | **C** | **D** |
| Linear | 97.1385 | 97.0106 | 97.4772 | 97.9619 |
| Nonlinear 2—3 | 2.0038 | 2.1600 | 1.7366 | 1.9275 |
| Nonlinear 4—5 | 0.8578 | 0.8295 | 0.7862 | 0.1106 |

**Table S2:** Parameter estimates for detection probability curves fitted on the protein group level for Datasets A–D visualized in Figure S3.

| | | **Detection probability curve** | |
|---|---|---|---|
| **Dataset** | **Fitted on** | $\beta_0$ | $\beta_1$ |
| A | Observed | -1.8880 | 0.8960 |
| | Underlying | -1.7533 | 0.8731 |
| B | Observed | -13.8707 | 1.2662 |
| | Underlying | -12.9501 | 1.1976 |
| C | Observed | -11.9163 | 0.5480 |
| | Underlying | -11.9005 | 0.5476 |
| D | Observed | -11.9030 | 0.7933 |
| | Underlying | -10.8627 | 0.7765 |

**Table S3:** Parameter estimates for detection probability curves fitted on Datasets E and F in Figures S4f–g and S5f–g.

| | | **Detection probability curve** | | |
|---|---|---|---|---|
| **Dataset** | **Level of quantification** | **Fitted on** | $\beta_0$ | $\beta_1$ |
| E | Precursor | Observed | -7.8223 | 0.5197 |
| | | Underlying | -7.7910 | 0.5184 |
| | Protein group | Observed | -7.1560 | 0.5761 |
| | | Underlying | -7.0890 | 0.5729 |
| F | Precursor | Observed | -12.5691 | 0.6063 |
| | | Underlying | -12.4444 | 0.6015 |
| | Protein group | Observed | -17.2926 | 0.8356 |
| | | Underlying | -17.1109 | 0.8295 |
